# A genetically-encoded cysteine biosensor to monitor cysteine dynamics across life domains

**DOI:** 10.64898/2026.01.27.698081

**Authors:** Brandán Pedre, Alexandre Deschamps, Charlotte Felten, Pauline Leverrier, Jean-François Collet, Bruno André, Peter Dedecker

**Affiliations:** Biochemistry, Molecular and Structural Biology Unit, Department of Chemistry, KU Leuven, Belgium; Molecular Physiology of the Cell Lab, Biopark - IBMM, Université Libre de Bruxelles, Gosselies, Belgium; WELBIO Department, WEL Research Institute, Avenue Pasteur 6, 1300, Wavre, Belgium; de Duve Institute, Université Catholique de Louvain, Avenue Hippocrate 75, 1200, Brussels, Belgium

## Abstract

Cysteine is a central metabolite in cellular redox regulation and iron-sulfur cluster assembly. Despite its critical role, monitoring cysteine dynamics in living systems has remained a challenge due to the lack of tools that avoid cysteine oxidation and/or do not destroy the cell in the process. Here, we report the development of Cystector (from <u>Cys</u>teine De<u>tector</u>), a genetically encoded, ratiometric green fluorescent biosensor for cysteine that exhibits an exceptional selectivity, minimal pH sensitivity in the physiological range, and a dynamic range of up to 4500%. Furthermore, the sensor retains functionality in the presence of physiological glutathione concentrations. We demonstrate the live-cell functionality of Cystector by monitoring intracellular and extracellular cysteine dynamics in different organisms. In *E. coli*, we show how cystine reduction in *Escherichia coli* is dependent on glutathione and glutaredoxins, and that the reduced cysteine is then exported into the extracellular environment. In yeast, we demonstrate how energy metabolism and oxidative stress determine cysteine homeostasis. In mammalian cells, we show how Cystector effectively monitors cysteine depletion in response to treatments such as H_2_O_2_, erastin2, and glutamate. Finally, we demonstrate, via a mitochondrially targeted variant, that Cystector can be used to monitor subcellular cysteine dynamics. These results together establish Cystector as a robust tool to unravel cysteine metabolism and transport in live cells across life domains.

## 1. Introduction

Besides serving as a proteogenic amino acid, cysteine is the limiting component in the synthesis of important non proteogenic biomolecules. This includes molecules such as glutathione, the main small thiol storage component involved in oxidative stress protection and xenobiotic resistance, and coenzyme A, a key cofactor in the oxidation of pyruvate in the citric acid cycle and the synthesis and oxidation of fatty acids. Cysteine is also the required sulfur donor for the formation of iron-sulfur clusters in proteins, which are essential to cellular processes such as the mitochondrial electron transport chain or DNA replication and repair (Lill and Freibert, 2020). Cells keep a delicate balance of cysteine concentrations, as an excess of cysteine is toxic and can be lethal (Pedre et al., 2021), with cells of different life origins having different strategies to cope with such excess. At the same time, insufficient cysteine supply in mammalian cells triggers cell death via ferroptosis (Dixon et al., 2012), with some tumor types being ‘addicted’ to sustained cysteine supply for their survival (Badgley et al., 2020; Cramer et al., 2017; Upadhyayula et al., 2023).

Because cysteine is actively exchanged between subcellular compartments and consumed by redox- and metabolism-linked processes, its intracellular concentration is dynamically regulated in space and time, as evidenced by lysosomal cyst(e)ine storage and mobilization (Adelmann et al., 2020; Kalatzis et al., 2001; Pisoni et al., 1990), mitochondrial cysteine maintenance under limiting cysteine conditions (Ward et al., 2024), or the control of cytoplasmic/cytosolic cysteine concentrations (Korshunov et al., 2020; Stipanuk et al., 2009). Unfortunately, current tools do not allow to effectively follow cysteine metabolism and transport in the native context of the cell, thus limiting our understanding on how cysteine spatiotemporal dynamics are regulated in the cell. Typical LC/MS-based pipelines to measure metabolite content do not work for cysteine, as the required cell lysis/tissue homogenization procedures oxidize the reduced cysteine pool. Pre-labelling cysteine with alkylating agents before metabolite extraction does not work either because the alkylation distorts the cysteine content (Bogdándi et al., 2019). Chemical fluorescent probes that measure cysteine are available but work via covalent modification, thus perturbing cysteine metabolism and preventing the measurement of cysteine dynamics (Jing et al., 2021). While two genetically-encoded fluorescent biosensors for free cysteine have been recently created, one functions by consuming cysteine and producing hydrogen sulfide, a toxic molecule (Caubrière et al., 2025), and the other has cross-reactivity to typical intracellular glutathione concentrations (Abrams et al., 2025), thus preventing a reliable monitoring of intracellular cysteine dynamics. Despite the progress in monitoring free cysteine, we still lack a tool that selectively and non-invasively monitors cysteine dynamics.

To fill the current technical gap, we herein have developed a genetically-encoded cysteine biosensor that is ratiometric, extremely selective to cysteine and pH resistant, that we named Cystector (from <u>Cyst</u>eine det<u>ector</u>). We demonstrate the utility of Cystector to monitor cysteine dynamics in *E. coli*, yeast and mammalian cells.

## 2. Design

To engineer Cystector, we first designed 25 chimeric proteins by inserting circularly permuted superfolder Venus (cpSFVenus from iGluSNFR3 v82 (Aggarwal et al., 2023)) into the *Neisseria gonorrhoeae* periplasmic cysteine solute receptor (Ngo2014) of the cysteine ABC transporter. The obtained crystal structure of this receptor naturally contains cysteine in its binding pocket and it was shown to specifically bind to cysteine rather than the structurally similar amino acid serine (Bulut et al., 2012). Among these Ngo2014-cpSFVenus chimeras, the cpSFVenus insertion after Ngo2014 A170 showed a 50% decrease in fluorescence intensity (excitation at 495nm) upon the addition of 2mM cysteine (Fig. S1A). We then randomized the composition of the linker residues connecting the split Ngo2014 with cpSFVenus. A linker variant containing one mutation at linker 1 (LSN to LVN) and two mutations at linker 2 (NNP to NHR) showed a 199% increase of fluorescence intensity upon cysteine addition (Fig. S1B). We then introduced three targeted mutations into cpSFVenus (I3R, L143F, G162S in cpSFVenus numbering) which made the biosensor dual-excitation, single emission ratiometric, with a 620% increase in the fluorescence ratio (R495/415) upon cysteine addition (Fig. S1C). To further increase the dynamic range, we applied two different strategies: one increased the linker 2 length and the other randomized a non-conserved Ngo2014 region located 3 residues after linker 2 (K174, S175, H176 in Ngo2014 numbering). Both strategies led to variants with increased dynamic ranges, ranging from 1500% to 2000% (Fig. S1D). However, the fluorescence ratio in all these variants was affected by pH changes typically found in the cytosol and mitochondrial matrix (7-8, Fig. S1E). In the final optimisation round, we randomized, via multisite-saturation mutagenesis, several positions within the fluorescent protein. This process obtained a 5-mutation variant (H21A, Y56T, V165I, Q166L, L211V, cpSFVenus numbering) that, when introduced in our previous variants, blue-shifted the maximum excitation wavelengths (400nm vs. 415nm, 470nm vs. 495nm), largely reduced the fluorescence ratio dependence on pH changes between 7 and 8, and increased the fluorescence ratio dynamic range (Fig. S1F). Considering the maximum fluorescence ratio change, robustness to pH changes and affinity/operating range within the reported intracellular cysteine levels (30-1000μM (Pedre et al., 2021)), we chose the variant with an additional residue at linker 2 (NHRF), denoted Cystector, for further characterization and testing in living cells.

## 3. Results

### A dual excitation ratiometric, cysteine specific, and pH resistant cysteine biosensor

Cystector has two excitation peaks around 400nm and 470nm that evolve in opposite directions in response to increasing cysteine concentrations (Fig. 1B). Both excitation at 400 and 470nm give rise to an emission peak in between 505 and 515nm (Fig. 1C-D). Upon binding to 2mM cysteine, Cystector showed a 257% decrease and an 832% increase in fluorescence intensity when excited at 400nm and 470nm, respectively, leading to a 4500% ratiometric fluorescence change. In terms of pH, fluorescence ratios both in absence of cysteine and with 2mM cysteine do not show major changes between pH 6.8-8 (Fig. 1E), indicating that this biosensor does not require a pH control within this pH range.

**Figure 1.**
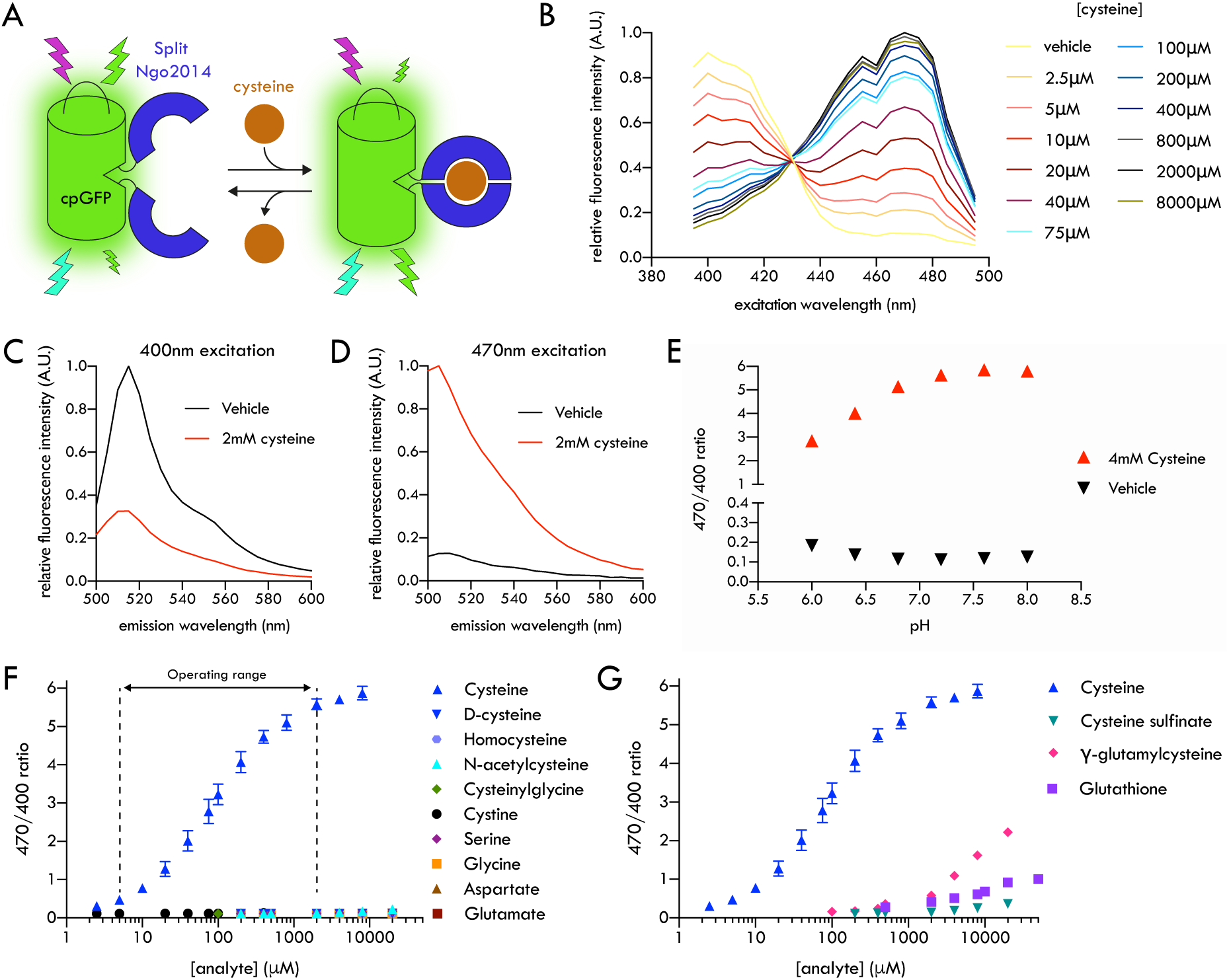
Characteristics of the cysteine biosensor Cystector. (A) Schematic composition and working mechanism of Cystector (B) Excitation scan (515nm emission, 20nm bandwidth) of purified Cystector under different cysteine concentrations (C) Emission spectra following excitation at 400nm in the absence and presence of 2mM cysteine (D) Emission spectra following excitation at or 470 nm in the absence and presence of 2mM cysteine (E) Ratio between excitation at 470nm and 400nm (R470/400, emission at 515nm emission, 20nm bandwidth) as a function of pH in the absence of cysteine and in the presence of 2 mM cysteine (F) R470/400-associated cysteine titration curve (K_d_=79.4μM, operating range 5-2000μM) and titration of structurally similar compounds: D-cysteine, serine, homocysteine, cysteinylglycine, N-acetylcysteine, and cystine. Data are shown as mean ± SEM (n=3 experimental replicates) (G) R470/400-associated response of Cystector to cysteinesulfinate, γ-glutamylcysteine, and glutathione. Data are shown as mean ± SEM (n=3 experimental replicates)

A titration with cysteine showed that Cystector has an apparent dissociation constant (K_d_) of 79.4μM at pH7.4, leading to an estimated operating range between 5μM and 2000μM (Fig. 1F). This operating range aligns well with intracellullar cysteine concentrations previously estimated to be in between 30μM and 1000μM (Pedre et al., 2021). Titration with other molecules demonstrated the selectiveness of Cystector to cysteine, as it showed no fluorescence response to structurally similar compounds such as D-cysteine, serine, homocysteine, cysteinylglycine, N-acetylcysteine or cystine (Fig. 1F). Some residual response was observed with cysteine sulfinate (Fig. 1G) and some cross reactivity was observed with γ-glutamylcysteine and glutathione (Fig. 1G). Among these, γ-glutamylcysteine triggered the strongest response, though the corresponding concentrations (≥2000μM) are ∼200-fold above the reported intracellular concentrations (∼5-10μM, (Mårtensson, 1987)). In the case of glutathione, the response of 5-10mM glutathione, considered the usual cytosolic concentration (Jiang et al., 2017), is equivalent to the response of 10μM cysteine (Fig. 1G). This cross-reactivity is over 10-fold less pronounced than that of a recent FRET-based cysteine biosensor, whose response to 10mM glutathione is greater than that seen for 100μM cysteine (Abrams et al., 2025).

To assess the potential interference of glutathione *in cellulo*, we repeated the cysteine titrations in presence of different glutathione concentrations. We observed that the presence of 6-20mM glutathione slightly reduces Cystector apparent binding affinity (79.4μM to 133.6μM in presence of 10mM glutathione), due to a reduced dynamic range in the lowest cysteine concentrations (Fig. S2A). Nevertheless, even when 10mM or 20mM glutathione is present, Cystector significantly detects 5 or 10μM cysteine, respectively (Fig. S2B-C). Given Cystector’s extreme selectivity to cysteine, resistance to pH changes, and excellent dynamic range, we decided to move on the testing into living systems.

### *E. coli* reduces cystine using glutathione and glutaredoxins and exports this cysteine into the extracellular environment

We first used Cystector in the *E. coli* BL21(DE3) strain to investigate cysteine import, as well as cystine import and its subsequent reduction to cysteine. We monitored both intracellular and extracellular cysteine levels by expressing Cystector within *E. coli* or by adding the purified Cystector to *E. coli* cells expressing a dark Cystector variant, respectively (Fig. 2A). Both cysteine and cystine addition led to an increase of the intracellular R470/400, indicating cysteine import or, in the case of cystine, its import followed by reduction to cysteine (Fig. 2B). We observed a faster R470/400 increase upon the addition of cystine, even though cystine needs to be reduced to be detected by Cystector (Fig. 2B). This faster cystine uptake has been previously reported and explained by the presence of dedicated cystine importers (Chonoles Imlay et al., 2015; Ohtsu et al., 2015) and the lack of a dedicated cysteine importer (Zhou and Imlay, 2020).

**Figure 2.**
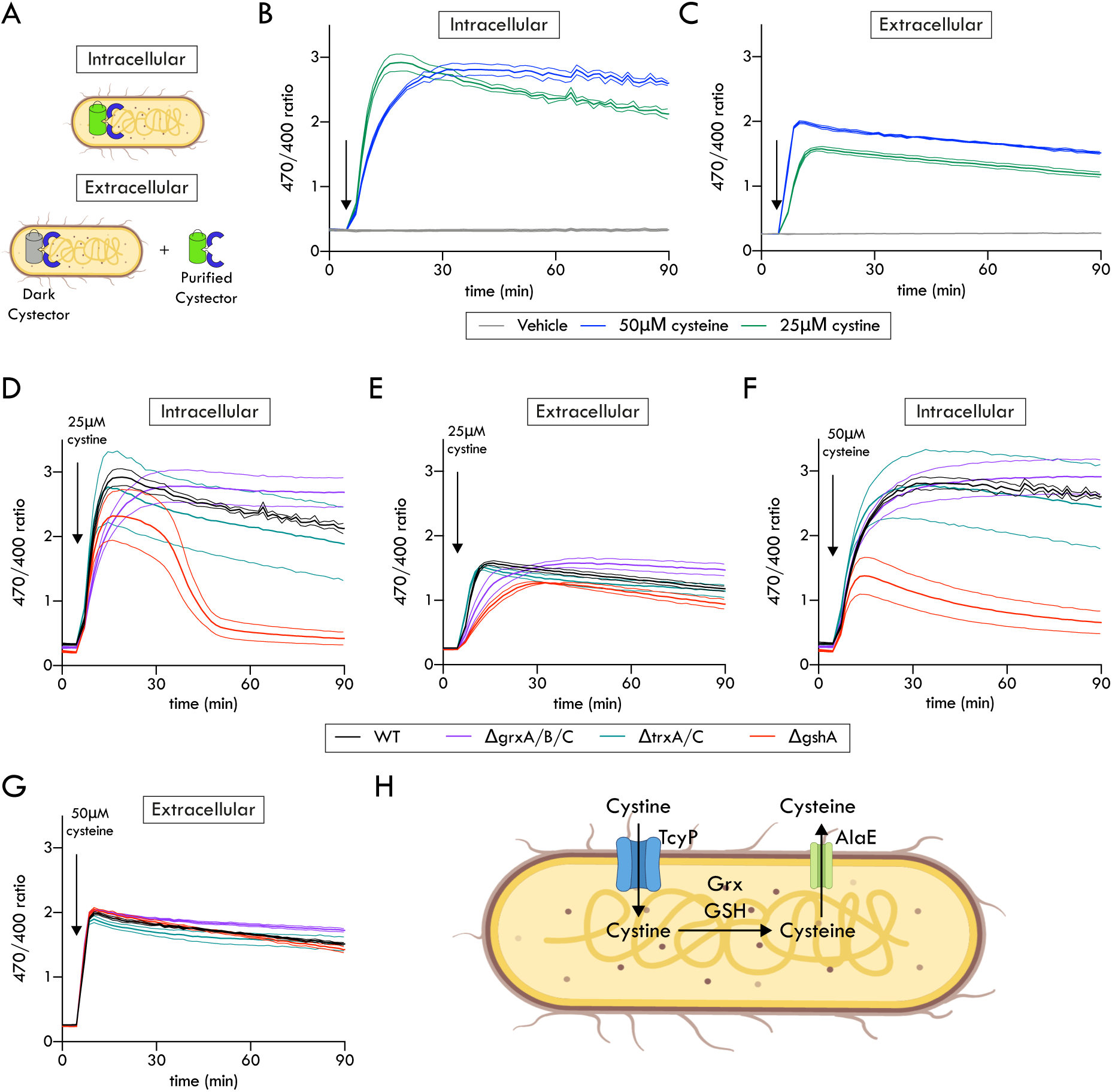
Monitoring cysteine and cystine uptake, reduction, and export in *E. coli*. (A) Experimental scheme for real-time monitoring of intracellular (using cytoplasm-expressed Cystector) and extracellular (by adding purified Cystector to the culture) cysteine levels (B) Intracellular Cystector R470/400 responses following addition of cysteine or cystine (C)Extracellular Cystector R470/400 responses following addition of cysteine or cystine (D) Intracellular Cystector R470/400 responses to cystine addition in deletion strains lacking thioredoxins (ΔtrxA/C), glutaredoxins (ΔgrxA/B/C), or γ-glutamylcysteine ligase (ΔgshA) (E) Extracellular Cystector R470/400 responses to cystine addition in deletion strains lacking thioredoxins (ΔtrxA/C), glutaredoxins (ΔgrxA/B/C), or γ-glutamylcysteine ligase (ΔgshA) (F) Intracellular Cystector R470/400 responses to cysteine addition in deletion strains lacking thioredoxins (ΔtrxA/C), glutaredoxins (ΔgrxA/B/C), or γ-glutamylcysteine ligase (ΔgshA) (G) Extracellular Cystector R470/400 responses to cysteine addition in deletion strains lacking thioredoxins (ΔtrxA/C), glutaredoxins (ΔgrxA/B/C), or γ-glutamylcysteine ligase (ΔgshA) (H) Proposed model of cystine dynamics in *E. coli*: upon import via TcyP, cystine is reduced in the cytosol, primarily by glutathione and glutaredoxins, and a fraction of this generated cysteine is exported via the amino acid exporter AlaE. Data are shown as mean ± SEM (n=3 experimental replicates) *E. coli* image and transporters were created with BioRender.com

Upon the addition of cystine, we also observed an increase of the extracellular Cystector R470/400, indicating that part of this cysteine formed intracellularly is exported into the extracellular environment (Fig. 2B). This result agrees with previous cystine import studies that reported an increase of extracellular thiols upon cystine addition, which the authors assumed to be cysteine but its identity was not confirmed (Korshunov et al., 2020).

Using the same intracellular/extracellular Cystector conditions, we then investigated the role of the major reducing systems in the *E. coli* cystine to cysteine conversion. Three deletion strains were studied: (1) Δ*gshA* (γ-glutamate-cysteine ligase), an enzyme that catalyzes the first step of glutathione synthesis, whose absence results in a strain that lacks glutathione; (2) Δ*grxA/B/C* (glutaredoxins), enzymes that reduce glutathione-containing mixed disulfides; and (3) Δ*trxA/C* (thioredoxins), enzymes that typically reduce protein disulfides. The Δ*trxA/C* strain did not show any impaired cystine reduction, while the Δ*grxA/B/C* strain revealed slower kinetics of cysteine appearance (Fig. 2D). The Δ*gshA* strain also showed a delay in its cystine reducing capacity: most importantly, after reaching a maximum R470/400 15 minutes after the addition of cystine, R470/400 sharply fell, indicating a dissipation of most of the intracellular cysteine (Fig. 2D). These cystine reducing delay in the Δ*grxA/B/C* and Δ*gshA* strains translates into a delayed extracellular R470/400 increase, indicating delayed cysteine export and, in the Δ*gshA* strain, a reduced cysteine export as well (Fig. 2E). In the case of cysteine addition, we observed a much smaller intracellular R470/400 increase in the Δ*gshA* strain, suggesting that the absence of glutathione also affects the reduced cysteine import (Fig. 2F). No strain differences in the R470/400 responses of extracellular Cystector were observed upon the addition of cysteine (Fig. 2G).

These results, along with previous studies, suggest the following cystine route in *E. coli*: the imported cystine by the transporter TcyP (Chonoles Imlay et al., 2015) is then reduced to cysteine in *E. coli* cytoplasm by glutaredoxins, and especially glutathione. Once reduced, excess cysteine is then exported to the extracellular medium, mediated by the broad spectrum amino acid exporter AlaE (Korshunov et al., 2020) (Fig. 2H).

### Yeast cysteine homeostasis depends on metabolic state and oxidative stress conditions

We next expressed Cystector in *Saccharomyces cerevisiae* to assess whether it could report intracellular cysteine dynamics in a eukaryotic microbial system. We first examined whether yeast cysteine content and homeostasis were different in cells growing under fermentative or post-diauxic shift respiratory conditions. We observed that, in absence of any treatment, Cystector R470/400 was about 3-fold lower under respiratory than under fermentative conditions (Fig. 3A), indicating cytosolic cysteine depletion. To probe cysteine homeostasis under both metabolic conditions, we added cysteine twice and monitored the Cystector R470/400 response. In fermentative cells, each cysteine addition induced a rapid increase in R470/400 that decreased toward a slightly elevated baseline within ∼30 min, suggesting efficient metabolization or mobilization of excess cysteine (Fig. 3B). On the other hand, under respiratory conditions, the first cysteine addition triggered an R470/400 increase which, as compared to fermenting cells, was of lower amplitude and did not return toward baseline, while the second cysteine addition did not show any further R470/400 response (Fig. 3B).

**Figure 3.**
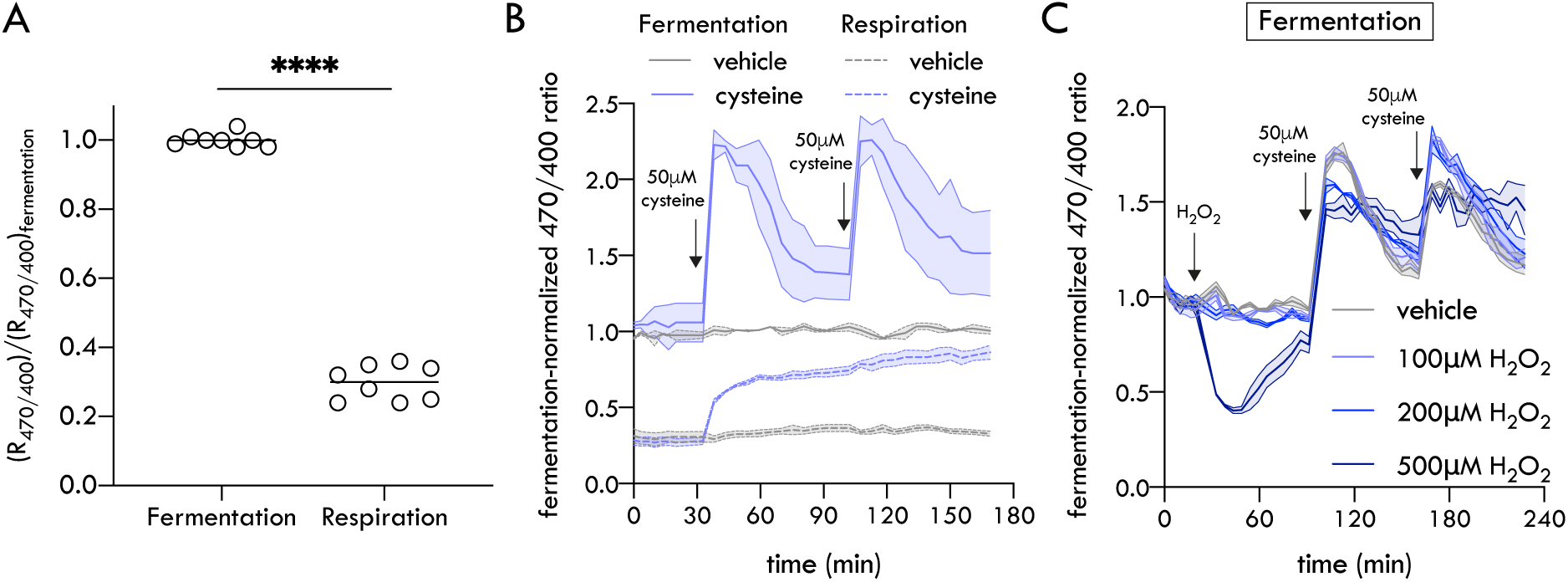
Differences in yeast cysteine homeostasis under different metabolic states and oxidative stress. (A)Steady-state Cystector R470/400 levels in yeast cells grown under fermentative vs. post-diauxic shift respiratory conditions (relative to fermentative conditions, data are shown as individual values + median, n=8 biological replicates, **** P < 0.0001 (unpaired two-tailed t-test with Welsch correction)). (B)Cystector R470/400 dynamics of yeast cells grown under fermentative and post-diauxic shift respiratory conditions upon repeated additions of cysteine. (C)Cystector R470/400 dynamics of yeast cells grown under fermentative conditions, treated with hydrogen peroxide and subsequent additions of cysteine. Data from B-D are shown as mean ± SEM (n≥2 biological replicates)

We then examined the Cystector response in yeast growing under fermentative conditions to the oxidant hydrogen peroxide (H_2_O_2_), which is known to oxidize the glutathione pool (Gutscher et al., 2008) and to deplete intracellular cysteine (Jing et al., 2021). Treatment with 100μM or 200μM H_2_O_2_ did not induce a noticeable decrease in the Cystector R470/400, whereas the exposure to 500μM H_2_O_2_ triggered a rapid R470/400 decrease that reached a plateau approximately 45 min after treatment, indicating cytosolic cysteine depletion (Fig. 3C). Following this initial decrease, the R470/400 slowly recovered toward its baseline value, indicating a gradual replenishment of intracellular cysteine (Fig. 3C). After these H_2_O_2_ treatments, we subsequently added cysteine twice and monitored the Cystector response. We observed that the H_2_O_2_ pretreatment altered cysteine homeostasis dynamics. In yeast cells pre-treated with low to intermediate H_2_O_2_ concentrations (100-200μM), cysteine addition elicited graded alterations in the Cystector R470/400 response. After the first cysteine addition, the R470/400 increases were of lower amplitude and the one at 200μM H_2_O_2_ returned to baseline more slowly. In both cases, the second cysteine addition produced a larger R470/400 increase than in untreated cells, indicating a partial relaxation of cysteine homeostasis (Fig. 3C). In contrast, in yeast cells pre-treated with 500μM H_2_O_2_, the characteristic rise-and-fall profile was no longer observed: the R470/400 remained elevated after the first cysteine addition, did not return toward baseline, and showed little further increase upon the second cysteine addition (Fig. 3C).

Together, these results suggest that, in glucose fermenting yeast, intracellular cysteine levels are higher though tightly regulated to prevent accumulation, as excess cysteine is toxic (Hughes et al., 2020). On the other hand, yeast under oxidative stress or post-diauxic shift respiratory conditions display lower cysteine levels and these levels are less dynamically regulated.

### Cystector effectively monitors intracellular cysteine depletion in mammalian cells

We also expressed Cystector in Cos7 cells, via transient transfection, to see if it could also detect cysteine dynamics in mammalian cells. Cystector was well expressed and its fluorescence intensity in plate reader measurements was higher than the autofluorescence of the cell culture medium (Fluorobrite DMEM +/− 2% fetal calf serum), enabling the simultaneous testing of multiple conditions.

We first tested what happens to the Cystector response upon treatment H_2_O_2_. The Cystector R470/400 decreased upon H_2_O_2_ treatment, indicating intracellular cysteine depletion, and this R470/400 decrease occurred in a H_2_O_2_ concentration-dependent manner (Fig. 4A). This decrease followed 2 phases: a “rapid” cysteine decrease, shared among the H_2_O_2_ treatments, and a slower but more pronounced decrease at higher H_2_O_2_ treatments (Fig. 4A). However, even this “rapid” decrease takes about 30 minutes post H_2_O_2_ treatment to reach a plateau (Fig. 4A). Subsequent experiments that used Cystector and the H_2_O_2_ biosensor HyPer7 in parallel wells (Pak et al., 2020) showed an immediate response of HyPer7 upon the H_2_O_2_ treatment, indicating that H_2_O_2_-induced disulfide formation precedes intracellular cysteine depletion (Fig S3). After the H_2_O_2_ treatment, we added cysteine to see whether the intracellular cysteine pool could be recovered: this happened with mild H_2_O_2_ doses (50-100μM) but not with a higher H_2_O_2_ concentration (200μM, Fig. 4A).

**Figure 4.**
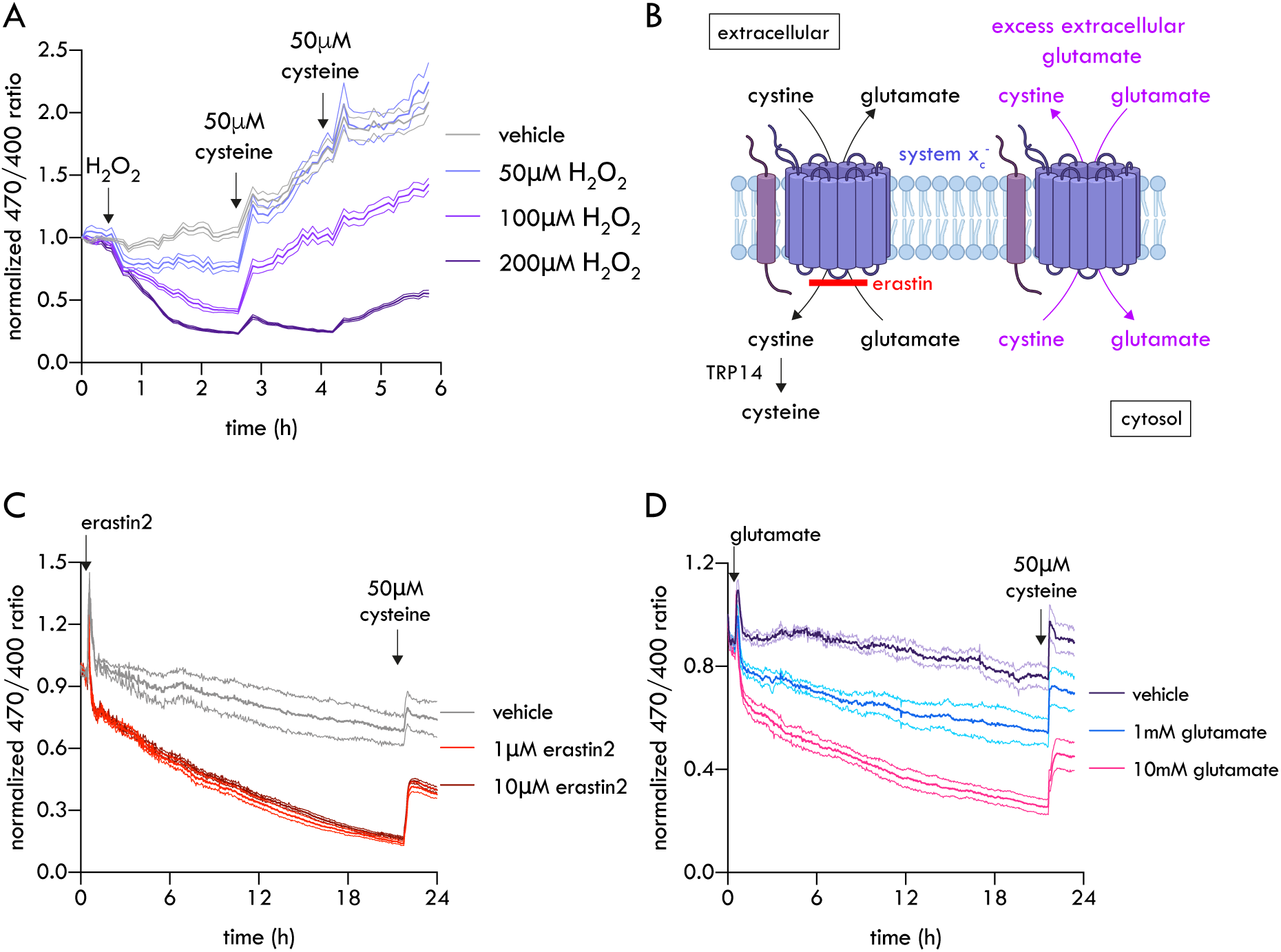
Cystector monitors cysteine depletion induced by oxidative stress and ferroptosis modulators in mammalian cells. (A)Cytosolic Cystector R470/400 responses in COS-7 cells following treatment with hydrogen peroxide (50-200μM) and subsequent treatments with 50μM cysteine (B)Schematic mechanism of action of treatments affecting cystine-glutamate antiporter system x_c_^-^, leading to intracellular cysteine depletion: erastin 2 blocks cystine import while glutamate drives cystine export (C)Cytosolic Cystector R470/400 responses in COS-7 cells following treatment with erastin2 (1 and 10μM) and the addition of 50μM cysteine the day after (D)Cytosolic Cystector R470/400 responses in COS-7 cells following treatment with glutamate (1 and 10mM) and the addition of 50μM cysteine the day after Data are shown as mean ± SEM (n≥3 biological replicates). Membrane and transporters were created with BioRender.com

We then tested what how the intracellular cysteine levels evolve upon treatment with the ferroptosis inducing compounds erastin and glutamate. These compounds reduce intracellular cysteine levels by acting on the cystine-glutamate antiporter system x_c_^-^, either by preventing cystine import (erastin (Sato et al., 2018; Yan et al., 2022)) or by turning x_c_^-^ into a cystine exporter (glutamate (Oka et al., 1993)) (Fig. 4B). However, we do not know how these treatments affect intracellular cysteine dynamics, as previous tools only allowed to determine cysteine uptake (Dixon et al., 2014) or endpoint intracellular cysteine measurements, usually the day after the treatment (Allen et al., 2023).

We observed, in Cystector-expressing cells, an immediate R470/400 decrease after erastin2 and glutamate treatments, indicating immediate cytosolic cysteine depletion (Fig. 4C-D). In the case of erastin2, both 1μM and 10μM treatments trigger a sudden R470/400 decrease 1h post treatment, followed by a more gradual decrease afterwards (Fig. 4C). In the case of glutamate, we observed the immediate R470/400 decrease being dependent on the applied concentration, followed by a gradual decrease less pronounced than the one of erastin2 (Fig. 4D). Finally, the R470/400 increase upon cysteine addition at the end of the experiment confirmed that Cystector remained functional and that the erastin2/glutamate treatments did not interfere with the reduced cysteine import (Fig. 4C-D).

### Mitochondrially-located Cystector reveals differences in yeast and mammalian cell cysteine homeostasis

Finally, we demonstrated that Cystector can be used to investigate subcellular cysteine distribution and dynamics in mammalian cells, by expressing a mitochondrial matrix-targeted variant in mammalian cell lines (Fig. 5A) and in yeast (Fig. 5B). In mammalian cells, under identical fluorescence measuring conditions at the plate reader, we observed that the steady-state R470/400 was higher in the mitochondrially-targeted Cystector than in the cytosolic version (Fig. 5C). Because Cystector R470/400 is not affected at this pH range (Fig. 1E), these results indicate that cysteine is more concentrated in the mitochondrial matrix than in the cytosol. On the other hand, we observed in yeast that the steady-state Cystector R470/400 was lower in mitochondria than in the cytosol under post-diauxic shift respiratory conditions, while no differences were observed under fermentative conditions (Fig. 5D). We then evaluated how the mitochondrial cysteine pool is affected by an external H_2_O_2_ dose and how it recovers upon the extracellular addition of cysteine. In the case of the mammalian cell line, all H_2_O_2_ treatments led to a faster R470/400 decline in cytosolic Cystector and, under mild treatments (50-100μM) the mitochondrial R470/400 showed a lower decrease to the one observed in the cytosol (Fig. 5E-G), indicating that the mitochondrial cysteine pool was less affected. In the case of the highest concentration, 200μM, both mitochondrial and cytosolic R470/400 decreased similarly (Fig. 5H). The addition of cysteine after the H_2_O_2_ treatment revealed different cysteine replenishment patterns: in absence of H_2_O_2_ or under a low H_2_O_2_ dose (50μM), similar R470/400 changes were observed upon the addition of cysteine, while in the highest H_2_O_2_ dose (200μM), the R470/400 increase was higher in mitochondrial Cystector (Fig. 5E-H). These results suggest that, under cysteine-limiting conditions, the recovery of the mitochondrial cysteine pool is prioritized over the cytosolic pool. In the case of yeast grown under fermentative conditions, however, the Cystector R470/400 changes upon H_2_O_2_ and the subsequent cysteine treatments were similar between the cytosolic and mitochondrial matrix-located biosensors (Fig. 5I-L). Together, these results show that Cystector reveals distinct, context-dependent modes of mitochondrial cysteine homeostasis in living cells.

**Figure 5.**
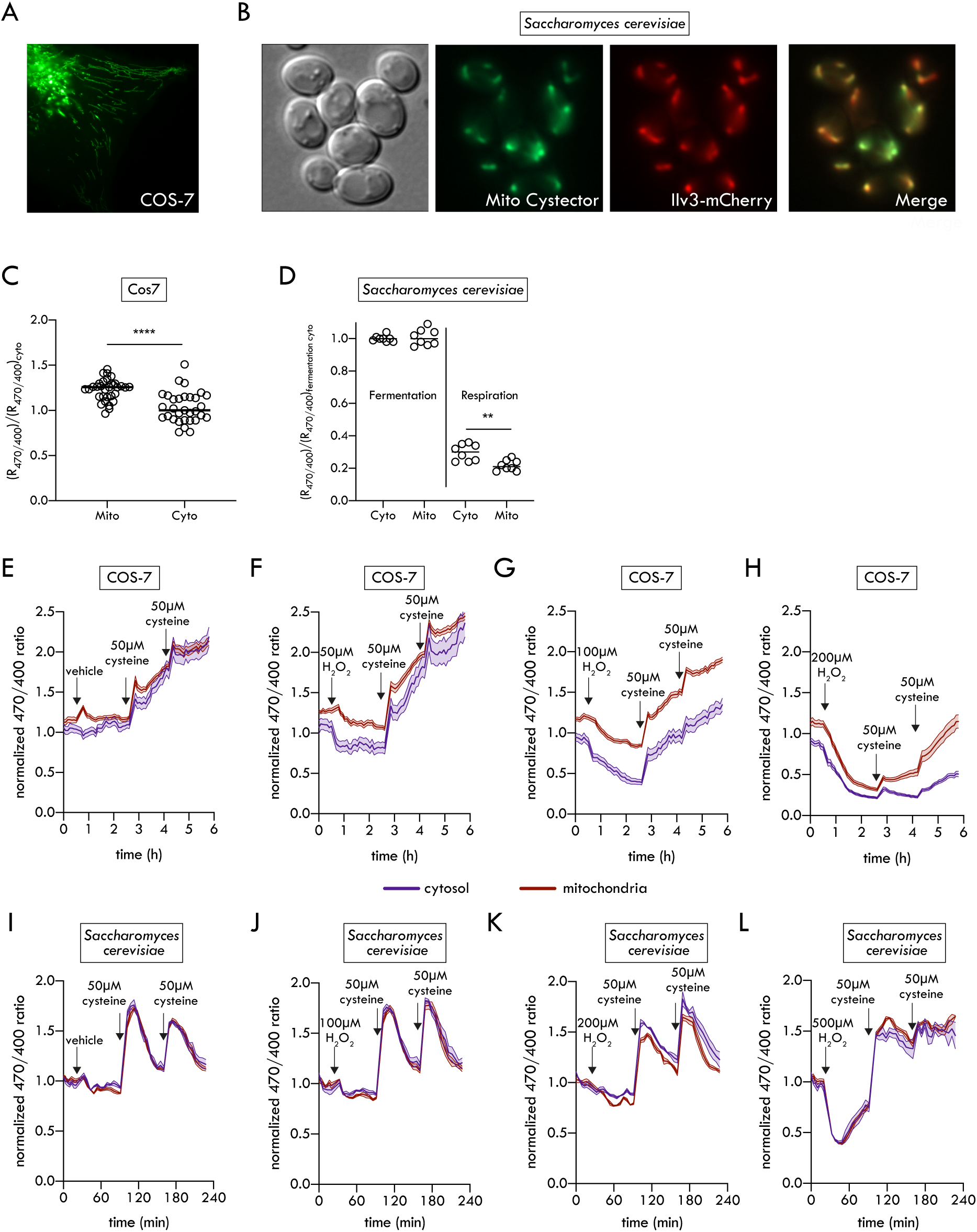
Differences in yeast and mammalian mitochondrial cysteine homeostasis. (A)Micrograph of mitochondrial matrix targeted Cystector upon excitation at 405nm. (B)Micrographs of mitochondrial matrix targeted Cystector (excitation at 470nm) co-expressed with chromosome-encoded mitochondrially targeted mCherry (excitation at 585nm) (C)Relative R470/400 between cytosolic and mitochondrial Cystector expressed in COS-7 cells (relative to cytosolic Cystector, n=31 biological replicates). **** P < 0.0001 (unpaired two-tailed t-test) (D)Relative R470/400 between cytosolic and mitochondrial Cystector expressed in yeast cells grown under fermentative or post-diauxic shift respiratory conditions (relative to cytosolic Cystector under fermentative conditions, data are shown as individual values + median, n=8 biological replicates). ** P < 0.01 (unpaired two-tailed t-test with Welsch correction). The dataset of cytosolic R470/400 values under fermentation conditions is identical to that presented in Fig. 3A. (E-H)Parallel monitoring of mitochondrial and cytosolic R470/400 changes (relative to cytosol) in COS-7 cells following treatment with hydrogen peroxide (vehicle -E-, 50μM -F-, 100μM -G-, 200μM -H-) and subsequent cysteine addition. Data are shown as mean ± SEM (n=6 biological replicates) (I-J)Parallel monitoring of mitochondrial and cytosolic R470/400 changes (relative to cytosol) in yeast grown under fermentative conditions following treatment with hydrogen peroxide (vehicle -I-, 100μM -J-, 200μM -K-, 500μM -L-) and subsequent cysteine addition. Data are shown as mean ± SEM (n=4 biological replicates). The data of cytosolic Cystector R470/400 are identical to those presented in Fig. 3D.

## 4. Discussion

For a biosensor to become a generalizable tool, it must be bright when expressed in cells, respond to the cognate analyte with high specificity and over its physiological concentration range, and not be affected by pH changes in the physiological pH range. Cystector meets these requirements and is in addition a ratiometric sensor with one of the highest dynamic ranges ever reported (4500%), if not the highest. This dynamic range is higher than recently developed biosensors such as the FiLa lactate sensor (1,500%-2,700%) (Li et al., 2023), iATPSnFR2 ATP sensor (∼1200%)(Marvin et al., 2024), or the redox biosensors roGFP2 (600-1200%)(Pedre, 2024), HyPer7 (500%)(Pak et al., 2020), SoNar (1500%)(Zhao et al., 2015) or iNap (900%) (Tao et al., 2017). Cystector also shows clear advantages over previous ratiometric cysteine biosensors. Unlike CyreB, which fuses roGFP2 with a cysteine desulfurase (Caubrière et al., 2025), Cystector does not require the desulfuration of cysteine (and therefore its consumption) to report changes in cysteine levels. Cystector also shows a considerably enhanced performance when compared to cpCys203, a recent FRET-based cysteine sensor that uses a homologue cysteine-binding protein and therefore has a similar sensing principle (Abrams et al., 2025). The key advantage of Cystector over cpCys203 is its higher selectivity towards cysteine over the highly abundant glutathione: while the FRET sensor responds more to 10mM glutathione than for 100μM cysteine, Cystector response to 10mM glutathione is equivalent to the one of 10μM cysteine, a 10-fold lower concentration. Given that intracellular cysteine levels in most cells range between 30μM and 240μM (Pedre et al., 2021), well below to the FRET sensor apparent binding affinity (313μM) (Abrams et al., 2025), and that glutathione concentrations are typically 5-10mM (Jiang et al., 2017), cpCys203 FRET-based are likely to be substantially confounded by glutathione, unlike those obtained with Cystector. Other advantages of Cystector over cpCys203 include its higher dynamic range (4500% vs 200%) and the use of a single fluorescent protein that prevents the artifacts inherent to FRET sensors, such as unequal fluorescent protein maturation or photodegradation.

We showed that Cystector can reveal new insights into cysteine metabolism and regulation through real-time monitoring of intracellular and extracellular cysteine dynamics in cells of different life domains. The studies in *E. coli* proved the hypothesis of Imlay and co-workers about the extracellular export of cysteine upon cystine reduction (Korshunov et al., 2020). We also demonstrated the involvement of glutaredoxins, but especially glutathione, in the cystine to cysteine reduction. Contrary to previous work in *E. coli* extracts, where Δ*grxA/B/C* was unable to reduce cystine (Chonoles Imlay et al., 2015), we observed that cystine reduction to cysteine in Δ*grxA/B/C* is slowed down but not abolished. It is thus possible that high glutathione levels and the presence of other reducing enzymes, such as glutathione disulfide reductase Gor, thioredoxins TrxA/TrxC or NrdH (Jordan et al., 1997) could partly take over the cystine to cysteine reduction role. An intriguing aspect is the fate of cystine in the Δ*gshA* strain, as it is reduced to cysteine but then disappears (Fig. 2D). The extracellular Cystector data indicate that the generated cysteine is not being exported (in fact, we observe less export -Fig. 2E-). Possible explanations include cysteine being transformed to other component (desulfhydration (Zhou and Imlay, 2022), racemization into D-cysteine), being mobilized to the periplasm, or cysteine being oxidized back to cystine because this strain has a lower reducing capacity. Recent work using roGFP2 in Δ*gshA* support the last hypothesis, as the cytosolic thiol-disulfide redox status in Δ*gshA* is much more oxidized (Caubrière et al., 2025; Knoke et al., 2025). As of why cysteine alone cannot compensate the role of glutathione to maintain a reducing environment, one explanation is related to the oxidation-prone nature of cysteine with Fe^3+^, which glutathione lacks (Park and Imlay, 2003).

The experiments in yeast illustrate how cysteine homeostasis is tightly and dynamically regulated by metabolic state and redox conditions. Under fermentative growth, yeast displayed higher basal cysteine levels and rapidly cleared excess cysteine following supplementation, indicating tight control of intracellular cysteine abundance. In contrast, under post-diauxic shift respiratory conditions, basal cysteine levels are lower, and cysteine accumulates after supplementation. These differences point to a profound remodelling of cysteine handling. The rapid cysteine clearance during fermentative growth is consistent with previous reports showing that the thiol group of supplemented cysteine is converted into H_2_S/sulfane sulfur in fermenting yeast (Winter et al., 2014), a process mediated by enzymes such as 3-mercaptopyruvate sulfurtransferase (TUM1 in yeast, (Huang et al., 2016)) and potentially by enzymes of the transulfuration pathway (Filipovic et al., 2018) or cysteinyl-tRNA synthetase CRS1(Nishimura et al., 2024), although direct evidence for H_2_S/sulfane sulfur production upon cysteine addition by the transulfuration/CRS1 enzymes in yeast has not been yet demonstrated. The vacuole is also involved in the cysteine sulfur conversion into H_2_S/sulfane sulfur, as mutants affected in vacuole acidification, vacuolar transport or vesicle fusion show dramatically lower H_2_S/sulfane sulfur production from cysteine (Winter et al., 2014), this being potentially linked to impaired cysteine partitioning into the vacuole. Of note, active cysteine transport into the vacuole has been reported to avoid the toxicity associated with cytosolic cysteine accumulation, which limits intracellular iron availability through an oxidant-based mechanism (Hughes et al., 2020).

In the case of the respiratory-grown cells, the lower basal cysteine levels are in agreement with reported lower levels of amino acids that originate from precursors of the glycolytic pathway, including the cysteine precursor amino acid serine (Martínez-force and Benítez, 1992), and the reported toxicity of excess cysteine on oxygen consumption (Bhuvaneswaran et al., 1964). The lack of cysteine clearance suggests a reduced expression or activity of the above-mentioned cysteine-catabolizing enzymes. Based on the loss of the cysteine clearance capacity in fermentative-growing cells upon treatment with H_2_O_2_, the lack of cysteine clearance in respiratory-grown cells could be caused by an increase of H_2_O_2_ or other reactive oxygen species due to the increased respiratory activity of mitochondria. While we did not identify the specific cysteine-regulatory components affected by H_2_O_2_, TUM1 represents a plausible target, as the activity of the human orthologue MPST is inhibited by H_2_O_2_ (Nagahara and Katayama, 2005), and oxidation of a TUM1-roGFP2 fusion biosensor was observed upon H_2_O_2_ addition (Pedre et al., 2023), consistent with oxidative inactivation of TUM1.

We also demonstrated that Cystector can be used to monitor intracellular cysteine dynamics in mammalian cells, as shown by the Cystector R470/400 decrease to cysteine depletion treatments (H_2_O_2_, erastin2, glutamate) and its increase upon the extracellular addition of cysteine. In the case of H_2_O_2_, the observed cysteine depletion must be caused by an indirect mechanism, considering that H_2_O_2_ is a poor oxidant of the cysteine thiol group (2.9M^-1^s^-1^) (Winterbourn and Metodiewa, 1999), especially when compared with the thiols in dedicated thiol peroxidase systems (≥10^5^M^-1^s^-1^ (Stone, 2004; Zeida et al., 2019)). One possible explanation is that H_2_O_2_ indirectly depletes cysteine by promoting the oxidation of reduced glutathione (GSH) into glutathione disulfide (GSSG), as observed using the Grx1-roGFP2 biosensor (Gutscher et al., 2008)). This reduces the cellular GSH pool. Since GSH exerts feedback inhibition on its own biosynthetic pathway (Richman and Meister, 1975), its depletion relieves this inhibition, triggering a compensatory increase in GSH synthesis that consumes intracellular cysteine. Another possible explanation is that the oxidative conditions from the H_2_O_2_ treatment trigger the oxidation of cystine reductase TRP14 (Martí-Andrés et al., 2024), which is then unable to reduce cystine or takes electrons from cysteine to recover its reduced state. These explanations support the intracellular cysteine decline but not why this decline is slow. One possibility is that lysosomes buffer the cytosolic cysteine loss, delaying the onset of detectable depletion. Lysosomes constitute a major reservoir of cellular cyst(e)ine (Abu-Remaileh et al., 2017), incorporating roughly half of newly synthesized cysteine despite comprising only ∼4% of cell volume (Pisoni et al., 1990), and oxidative cues such as H₂O₂ can mobilize this pool for GSH biosynthesis (He et al., 2023). This buffering could explain why the Cystector response evolves slowly despite the rapid thiol oxidation detected by HyPer7.

Cystector also revealed that the ferroptosis inducers erastin2 and glutamate lower the intracellular cysteine levels. Although both treatments result in a sustained depletion of intracellular cysteine, the most pronounced decrease occurs within the first hour following erastin2 or glutamate addition. Previous studies in cystine-starved cells reported rapid cytosolic cysteine depletion with a half-life of approximately 22 minutes (Ward et al., 2024), yet the decline observed with Cystector was comparatively modest relative to the ∼85% cysteine loss in the first hour that was reported from LC-MS measurements. This difference likely indicates that cystine starvation constitutes a more severe perturbation of cellular cysteine homeostasis than pharmacological inhibition of system x_c_^-^ by erastin2 or glutamate, leading to a more rapid and extensive depletion of the intracellular cysteine pool.

The long-term cysteine depletion can be readily explained by the inhibition of cystine import (erastin) or promotion of cystine efflux (glutamate), which deprives the cell of its primary extracellular cysteine precursor. However, the rapid onset of depletion suggests the involvement of additional mechanisms. Recent work shows that system x_c_^-^ is also present in lysosomes and erastin inhibits its cystine efflux activity, which would restrict access to lysosomal cyst(e)ine stores (Zhou et al., 2025).

Finally, by targeting Cystector to the mitochondrial matrix, we demonstrated its use to monitor cysteine dynamics in subcellular compartments. We showed that mild H_2_O_2_ doses affect the mammalian mitochondrial cysteine pool less than the cytosolic one. While this sounds intuitive, since H_2_O_2_ would have to traverse 3 membranes to exert its effect at the mitochondrial matrix, thiol-based redox biosensors Grx1-roGFP2 and HyPer7 report the opposite effect, with H_2_O_2_ leading to a higher oxidation in the mitochondrial matrix than in the cytosol (Hoehne et al., 2022; Secilmis et al., 2021; Swain et al., 2016). This means that mitochondrial thiol redox status and cysteine content are not entirely coupled. In addition, the cysteine-induced recovery upon cysteine depletion with 200μM H_2_O_2_ indicates that, under conditions where cysteine levels are limited, mitochondria preferentially uptake this cysteine. This is not unexpected considering that, as mitochondria require the sulfur atom of cysteine to assemble iron-sulfur clusters (Lill and Freibert, 2020), present in mitochondrial and extramitochondrial proteins, it would make sense that cysteine is preferentially supplied to mitochondria. On the other hand, yeast grown under fermentative conditions show no differences in cysteine concentration between the mitochondrial matrix and the cytosol or preferential accumulation of cysteine into the cytosol under oxidative stress. The lack of preferential mitochondrial cysteine accumulation is consistent with the reduced demand for mitochondrial iron-sulfur cluster biosynthesis under fermentative growth, when enzymes of the citric acid cycle and respiratory chain are largely dispensable. In both mammalian cells and yeast, the sudden increase of the mitochondrial cysteine levels upon the extracellular addition of cysteine suggests the presence of a cysteine importer, which has been already proposed but not identified yet.

## 5. Limitations of the study

This study uses 3 model living systems: *E. coli, S. cerevisiae*, and COS-7 mammalian cells, to demonstrate the cross-domain applicability of Cystector. However, it is possible that the expression of Cystector in other living systems (other bacteria, zebrafish, plants) needs to be optimized with codon optimized variants. In those cases, the authors are happy to discuss with those interested in Cystector how its engineering can improve the expression at the host organism. Fluorescence measurements using 400nm excitation can be challenging due to high background autofluorescence from cells and culture media in this spectral region, as well as reduced tissue or sample penetrance of violet light. In such cases, we recommend fusing Cystector to a bright red fluorescent protein, such as mScarlet3 (Gadella et al., 2023)) and quantifying cysteine dynamics by calculating the ratio between green fluorescence upon 470 nm excitation and red fluorescence emission.

## 6. Methods

### 6.1 Reagents

L-cysteine-HCl, L-cystine, L-cysteine sulfinate, γ-glutamylcysteine, cysteinylglycine, homocysteine glycine, N-acetylcysteine, serine, aspartate, glutamate, hydrogen peroxide (H_2_O_2_), tryptone, Tris, dimethylsulfoxide ampicillin, kanamycin and ethylenediaminetetraacetic acid (EDTA) were purchased from Sigma-Aldrich. Glutathione (reduced form, 99%) was purchased from Roth. Imidazole, sodium chloride, sodium hydroxide, hydrochloric acid sodium ortophosphate and mammalian cell culture components (DMEM, Fluorobrite DMEM, trypsin, glutaMAX, fetal calf serum (FCS)) were purchased from Thermo Scientific. Erastin2 was purchased from Cayman Chemical. Di-hydrogen sodium phosphate was purchased from Chem-Lab. Yeast extract was purchased from Fisher Scientific. Lactose was purchased from UCB. Glucose was purchased from Acros Organics. HEPES was purchased from VWR. A 1mM L-cystine stock solution was prepared by dissolving it in 100mM sodium hydroxide and pH was subsequently adjusted to 7.4 with hydrochloric acid. A 10mM erastin2 stock solution was prepared by dissolving it in dimethylsulfoxide.

### 6.2 Plasmid construction and mutagenesis

JM109(DE3) was the host strain used throughout the Cystector engineering process. The components of the 1st engineering generation (Ngo2014 8-170, cpSFVenus from iGluSNFR3 v82 (Aggarwal et al., 2023), Ngo2014 171-284) were amplified by PCR using high-fidelity DNA polymerase Q5 (New England Biolabs), and contained overhanging fragments that enabled the assembly of these components together and with vector pRSET-B (BamHI/EcoRI digested) using HiFi DNA Assembly (New England Biolabs). In the 2nd generation, linker libraries were created via HiFi DNA Assembly of two components: (i) all the plasmid except the linker residues and cpSFVenus and (ii) cpSFVenus and the linker residues, the latter randomized by introducing degenerate NNK codons (or MNN at the reverse primer) at positions 2 and 3 in both linkers. The 3rd generation used three sequential rounds of site-directed mutagenesis, which followed the specifications of QuikChange - Site-Directed Mutagenesis Kits (Agilent) but using Q5 high-fidelity DNA polymerase (New England Biolabs) for PCR amplification, and FD DpnI (Thermo Scientific) for template digestion. The 4th generation used site-saturation mutagenesis with degenerate NNK codons (or MNN at the reverse primer) to (i) introduce an extra linker residue at the C-terminus of linker 2 or (ii) to randomize the amino acid composition of Ngo2014 K174-H176 region. Similar amplification specifications as with site-directed mutagenesis (Q5 DNA polymerase, FD DpnI) were used. The 5th generation created site-saturated mutant libraries within cpSFVenus at positions 21, 56, 165, 166 and 211, via HiFi DNA Assembly of four PCR-amplified fragments: (i) all the plasmid outside cpSFVenus positions 21 and 211; (ii) cpSFVenus positions 21 and 56, both primers containing NNK/MNN codons at these positions; (iii) cpSFVenus positions 57 and 164; and (iv) cpSFVenus positions 165 and 211, with fw primer containing NNK codons at positions 165 and 166, and MNN codon at position 211.

A pRSET-B dark Cystector was generated by introducing a chromophore-disabling Y-F mutation via mutagenesis PCR.

For yeast expression, a codon-optimized version of Cystector was ordered (Integrated DNA Technologies), PCR amplified and assembled into p416TEF1 (XbaI/HindIII digested) (Mumberg et al., 1995) using HiFi DNA Assembly. For expression into yeast mitochondria, p416TEFSu9DAAO(Pedre et al., 2023), which contains F_0_-ATPase subunit 9 (Su9) from *Neurospora crassa* -mitochondrial targetting sequence- and D-amino acid oxidase (DAAO) from *Rhodotorula gracilis*, was NcoI/HindIII digested to release DAAO and clone the codon-optimized Cystector C-terminally of Su9 via HiFi

For mammalian cell expression, the final Cystector from the *E. coli* engineering (minus the His-tag) was subcloned into a pCDNA3.1 vector (HindIII/XbaI digested) via HiFi DNA assembly. For expression into mammalian cell mitochondria, the pCDNA3.1 Cystector (HindIII digested) was C-terminally fused to a 2xCOXVIII mitochondrial targetting presequence via HiFi DNA assembly.

### 6.3 Library variant screening

Transformant colonies were visually inspected for green/yellow fluorescence upon illumination with violet and blue light. Bright variants were picked and grown in 600μL autoinduction media (20g/L tryptone, 5g/L yeast extract, 5g/L NaCl, 15.15g/L Na_2_HPO_4_x12H_2_O, 3g/L KH_2_PO_4_, 0.6% glycerol, 0.005g/L glucose, 0.02g/L lactose, 100μg/mL ampicillin) within a 96-deep well block (Thermo Scientific), and grown at 37°C overnight under shaking. Cultures were then spun down (4000rpm, 15min), washed once with 600μL assay buffer (200mM sodium phosphate buffer, pH7.4, 150mM NaCl and 0.5mM EDTA) and lysed with 100μL Bacterial protein extraction reagent B-PER (Thermo Scientific) plus vortexing. After lysis, 500 µL assay buffer were added to dilute the extract. The insoluble cell debris was removed by centrifugation (4000rpm, 30min) and 180μL of the cleared lysate were transferred to a 96-well plate. The plate was then incubated at 37°C for 15min in the plate reader (Tecan Spark) and 2 excitation scans were collected, before and after addition of 2mM cysteine (390-495nm, emission 535nm -generations 1 to 4-; 380-480nm, emission 515nm -generation 5-; 5nm excitation bandwidth, 5nm steps, 20nm emission bandwidth).

### 6.4 Recombinant protein expression and purification

The plasmid (pRSETb) containing His-tagged Cystector was transformed into *E. coli* JM109(DE3). A single colony was inoculated with 300mL of autoinduction media supplemented with 100 µg/mL ampicillin. The culture was shaken at 21°C (180-200 rpm) for 72 hours. The cells were then harvested and lysed with B-PER containing 5mM imidazole, 1X cOmplete™, Mini, EDTA-free Protease Inhibitor Cocktail (Roche). The resulting cellular debris was centrifuged for 20 minutes at 8500 rpm, and the clarified lysate was loaded onto 5mL Pierce polypropylene columns (Thermo Fisher Scientific) containing 2mL of a Ni-NTA agarose suspension (Qiagen). The column was then washed with 20mL of 100 mM Tris-HCl pH 7.4, 300mM NaCl and 20 mM imidazole, and eluted with the same buffer but containing stepwise increases of imidazole concentration (50, 100 and 250mM). The eluted fractions were analysed for purity on SDS-PAGE: the fractions considered pure enough were then pooled and concentrated to ∼5mL using a Vivaspin 6 10,000 MWCO PES (Sartorius) membrane ultrafiltration tube. The excess imidazole was removed via buffer exchange using two PD-10 desalting columns (Cytiva) pre-equilibrated with 15mM Tris-HCl pH7.4, 150mM NaCl, 1mM EDTA. The purified protein was stored at 4°C.

### 6.5 In vitro fluorescence spectroscopy experiments

All measurements were performed at 37°C in Tecan Spark plate reader using Greiner 96 Flat Black Chimney Well Plates and a fixed gain. Cystector was diluted in assay buffer to a final concentration of 100 nM in a final reaction volume of 200μL per well (180μL of diluted Cystector plus 20μL of assay buffer or 10X concentrated analyte solution -dissolved in assay buffer-). Excitation scans were recorded from 390 to 495nm (5nm steps, 5nm bandwidth), with emission set at 515nm (20nm bandwidth). Emission scans were recorded from 500 to 600nm (5nm steps, 5nm bandwidth) after excitation at 400nm or 470nm (20nm bandwidth in both). For pH-dependence measurements, purified Cystector was diluted in assay buffer with varying proportions of NaH_2_PO_4_/Na_2_HPO_4_ to reach pH 6, 6.4, 6.8, 7.2, 7.6 and 8. For glutathione-competition measurements, 160μL of diluted Cystector were mixed with 20μL of a 10X-concentrated glutathione solution, followed by addition of 20μL of a 10X-concentrated cysteine solution.

The determination of the apparent dissociation constant (K_d_) in the cysteine titration assay was done by fitting the data to a one site-total binding equation (Y=B_max_*X/(K_d_+X) + NS*X + Background) in GraphPad Prism, where B_max_ stands for maximum ratio and NS for non-specific binding.

### 6.6 *E. coli* cysteine-based fluorescence measurements

*E. coli* experiments were performed in strain BL21(DE3). Mutant strains Δ*gshA*, Δ*grxA/B/C* and Δ*trxA/C* were created as previously described (Caubrière et al., 2025) Briefly, the corresponding Keio kan^R^ alleles were introduced by P1 phage transduction, kanamycin insertions were verified by PCR, and kan^R^ cassettes were excised by transformation with pCP20 plasmid when additional deletions were required. All strains were made chemically competent by the Inoue method (Inoue et al., 1990) and transformed with Cystector/dark Cystector-containing pRSET-B via heat shock.

Colonies of BL21(DE3) Cystector and dark Cystector transformants were grown overnight at 37°C in autoinduction media. The culture OD_600nm_ was measured, the cells were harvested, washed once in PBS, and finally resuspended in PBS to reach a final OD_600nm_ of 7. The dark Cystector-containing suspension was divided in half and 100nM purified Cystector was added to the second half. The three suspensions (dark Cystector -background fluorescence control-, dark Cystector + purified Cystector, Cystector) were then dispensed onto a Greiner 96 Flat Black Chimney Well Plate (180μL/well) and preincubated at 37°C within the Tecan Spark plate reader for 20min before starting the measurement. Fluorescence intensity measurement cycles at 400nm and 470nm excitation (both at 20nm bandwidth, emission at 515nm, 20nm bandwidth, fixed gain) were taken every 1.5min. Baseline fluorescence measurements were acquired for 10min, followed by the addition of 20μL cysteine (500μM) or cystine (250μM).

### 6.7 *S. cerevisiae* cysteine-based fluorescence measurements

Experiments in yeast used strain BY4742 (*MATα his3Δ, leu2Δ, lys2Δ ura3Δ*), which was transformed with pCJ315 plasmid (to complement *his3Δ, leu2Δ, lys2Δ* auxotrophies (Saliba et al., 2018)) and with either p416TEF1 cytosolic or mitochondrially-located Cystector, using the lithium acetate method. Cells were grown at 29 °C in minimal medium buffered at pH 6.1(Jacobs et al., 1981), with glucose (0.003g/L) as carbon source and (NH_4_)_2_SO_4_ (10 mM) as nitrogen source. Prior to the assay, cells were harvested by filtration, washed, and resuspended in fresh medium kept at 29°C. Cell suspensions were adjusted to equivalent OD_600nm_ and dispensed into a black Greiner 24-well plate (reference: 662174). Fluorescence intensity measurement cycles at 400nm and 470nm excitation (both at 20nm bandwidth, emission at 511nm, 20nm bandwidth, fixed gain) were taken every 3min20s or 5min20s.

### 6.8 COS-7 cysteine-based fluorescence measurements

A T25 flask containing 10^6^ seeded COS-7 cells was transfected with 2.5μg pCDNA3.1 cyto Cystector, mito Cystector, or HyPer7, and 15μL of transfection reagent FuGene6 (Promega). The next day, the cells were trypsinized and seeded onto a Cellstar Greiner 96 μClear Black Chimney Well Plate (25000cells/well). Untransfected COS-7 cells were seeded in the same plate for background fluorescence substraction. For long term-experiments (erastin2/glutamate treatments), seeding was done in Fluorobrite DMEM with glutaMAX and 2% FCS, and media was not replaced for the measurement. For short-term experiments (H_2_O_2_), seeding was done in DMEM with glutaMAX and 10% FCS, and the day after the cells were washed with PBS and replaced with 200μL/well of Fluorobrite DMEM with glutaMAX.

Fluorescence measurements at the Tecan Spark plate reader were done at 37°C and 5% CO_2_. Fluorescence intensity measurement cycles at 400nm and 470nm excitation (both at 20nm bandwidth, emission at 515nm, 20nm bandwidth, 60% gain for 400nm and 40% gain for 470nm within the plate) were taken every 2.5-5min. Baseline fluorescence measurements were acquired for ∼15min, followed by the treatments (10X of H_2_O_2_, glutamate or cysteine solutions in PBS, 10X of erastin2 solution in fluorobrite DMEM + glutaMAX and 2%FCS).

### 6.9 Fluorescence microscopy measurements

Mito Cystector-transfected Cos7 cells were imaged on a Nikon Eclipse Ti-2 Inverted Microscope (Minato City, Japan) equipped with a 1.4 NA oil immersion objective (100× CFI Apochromat Total Internal Reflection Fluorescence) and a ZT405/488/561/640rpcv2 dichroic mirror with a ZET405/488/561/640 nm emission filter (both Chroma Technology, Bellows Falls, Vermont) in epi-illumination. Two separate lasers at 405 and 488 nm (Oxxius, Lannion, France), at 20% power, were used for excitation. Images were acquired on a PCO. edge 4.2 camera (PCO, Kelheim, Germany) with an exposure time of 50ms. Image analysis was performed using IGOR Pro(Wavemetrics).

Mito Cystector-transformed yeast cells (strain Σ1278b) that co-express the mitochondrial marker Ilv3-mCherry from the chromosome were imaged by epifluorescence microscopy on a Zeiss Axio Imager.M2 microscope, with a Colibri 2 LED light source. The images were acquired using a Plan-Apochromat 100x/1.4 NA oil immersion objective (Zeiss) and a Zeiss AxioCam MRm monochrome CCD camera. Cystector was illuminated at 470nm with GFP filter set 38, dichroic at 495 nm and emission at 525/50nm, while mCherry was illuminated at 555nm with mCherry filter set 45, dichroic at 585 nm and emission at 630/75nm.

## 7. Materials availability

The following plasmids generated in this study have been deposited to Addgene: pRSET-B-6xHisTag-Cystector (Cat# 250652), pCDNA3.1-Cystector (Cat# 250653), p416TEF1-cytoCystector (Cat# 250654), p416TEF1-Su9(mito)Cystector (Cat# 250655). Additional constructs generated in this study are available from the lead contacts with a completed materials transfer agreement.

## 8. Acknowledgements

Brandán Pedre thanks Jiashuo Zheng (Helmholtz Zentrum München) for having proposed the name Cystector for the cysteine biosensor after having met at the 2024 Gordon Research Conference on Thiol-Based Redox Regulation and Signaling. Brandán Pedre thanks Mariona Moranta Coll (IES Son Pacs) for her research project enabling the investigation of *E. coli* extracellular cysteine efflux using purified Cystector. Brandán Pedre thanks the Research Foundation-Flanders (FWO Vlaanderen) for his postdoctoral fellowship (grant number 1276324N). This work was supported by the Research Foundation-Flanders through grants G090819N and G010723N, and the KU Leuven via C14/22/088.

