## Supporting information for "A genetically-encoded cysteine biosensor to monitor cysteine dynamics across life domains"

#### Supplementary information

#### Figures S1-3

**Figure S1. Overview of the Cystector development process**
(A)Excitation scan (emission 535nm, 20nm bandwidth) of the initial Ngo2014-170–cpSFVenus insertion construct in absence or presence of 2mM cysteine
(B)Excitation scan (emission 535nm, 20nm bandwidth) of linker-optimized version (linker 1: LSN→LVN; linker 2: NNP→NHR) in absence or presence of 2mM cysteine
(C)Excitation scan (emission 535nm, 20nm bandwidth) of third-generation version with 3 point mutations (I3R, L143F, G162S, cpSFVenus numbering) in absence or presence of 2mM cysteine
(D)Excitation scans (emission 535nm, 20nm bandwidth) of fourth-generation variants generated via linker 2 editing or randomization of Ngo2014 K174, S175, H176 residues (Ngo2014 numbering) in absence or presence of 2mM cysteine
(E)Ratio between excitation at 495nm and 415nm (R495/415, emission at 535nm emission, 20nm bandwidth) as a function of pH in the absence of cysteine and in the presence of 2 mM cysteine
(F)Excitation scan (emission 515nm, 20nm bandwidth) of Cystector, fifth and final generation variant, generated via multiple mutations within cpSFVenus (H21A, Y56T, V165I, Q166L, L211V)

**Figure S2. Effect of glutathione on Cystector affinity and detection sensitivity.**
(A)Cysteine titration curves of Cystector in the absence or presence of different glutathione concentrations (6-20 mM).
(B)Cystector R470/400 response to different cysteine concentrations in presence of 10mM glutathione. ** $P < 0.01$ (unpaired two-tailed t-test)
(C)Cystector R470/400 response to different cysteine concentrations in presence of 20mM glutathione * $P < 0.05$ (unpaired two-tailed t-test)
Data are shown as mean ± SEM (n=3 experimental replicates)

**Figure S3. Comparative kinetics of hydrogen peroxide sensing and cysteine depletion in mammalian cells**

Parallel monitoring of intracellular hydrogen peroxide dynamics using HyPer7 and cysteine dynamics using Cystector in COS7 cells following repeated 50μM $H_2O_2$ treatments. Data are shown as mean ± SEM (n=3 experimental replicates)

### Figure S1

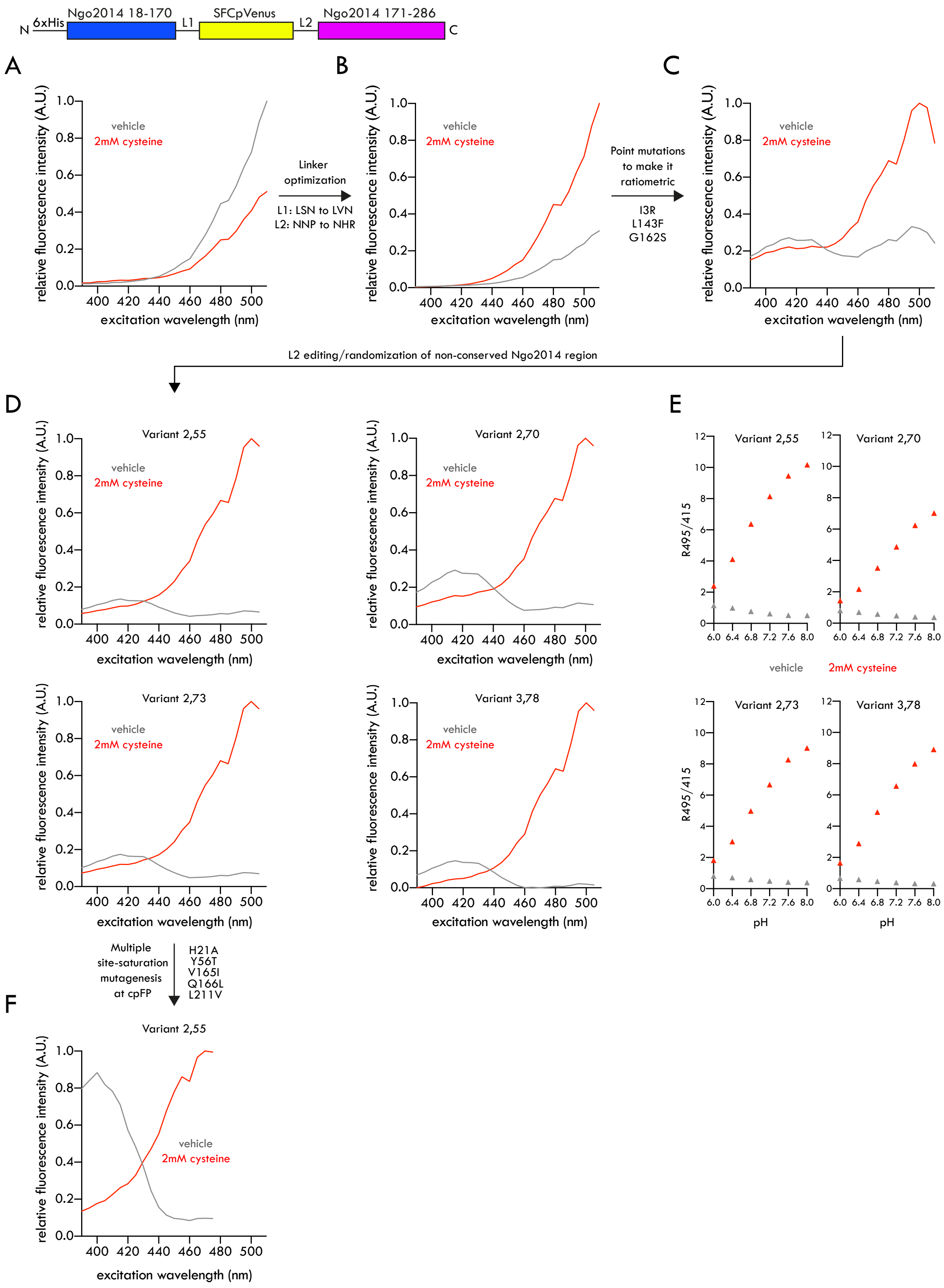
N 6xHis Ngo2014 18-170 L1 SFCpVenus L2 Ngo2014 171-286 C
A
B
C
D
E
F
relative fluorescence intensity (A.U.)
excitation wavelength (nm)
vehicle
2mM cysteine
Linker optimization
L1: LSN to LVN
L2: NNP to NHR
Point mutations to make it ratiometric
I3R
L143F
G162S
L2 editing/randomization of non-conserved Ngo2014 region
Variant 2,55
Variant 2,70
Variant 2,73
Variant 3,78
R495/415
pH
Multiple site-saturation mutagenesis at cpFP
H21A
Y56T
V165I
Q166L
L211V

#### Figure S2

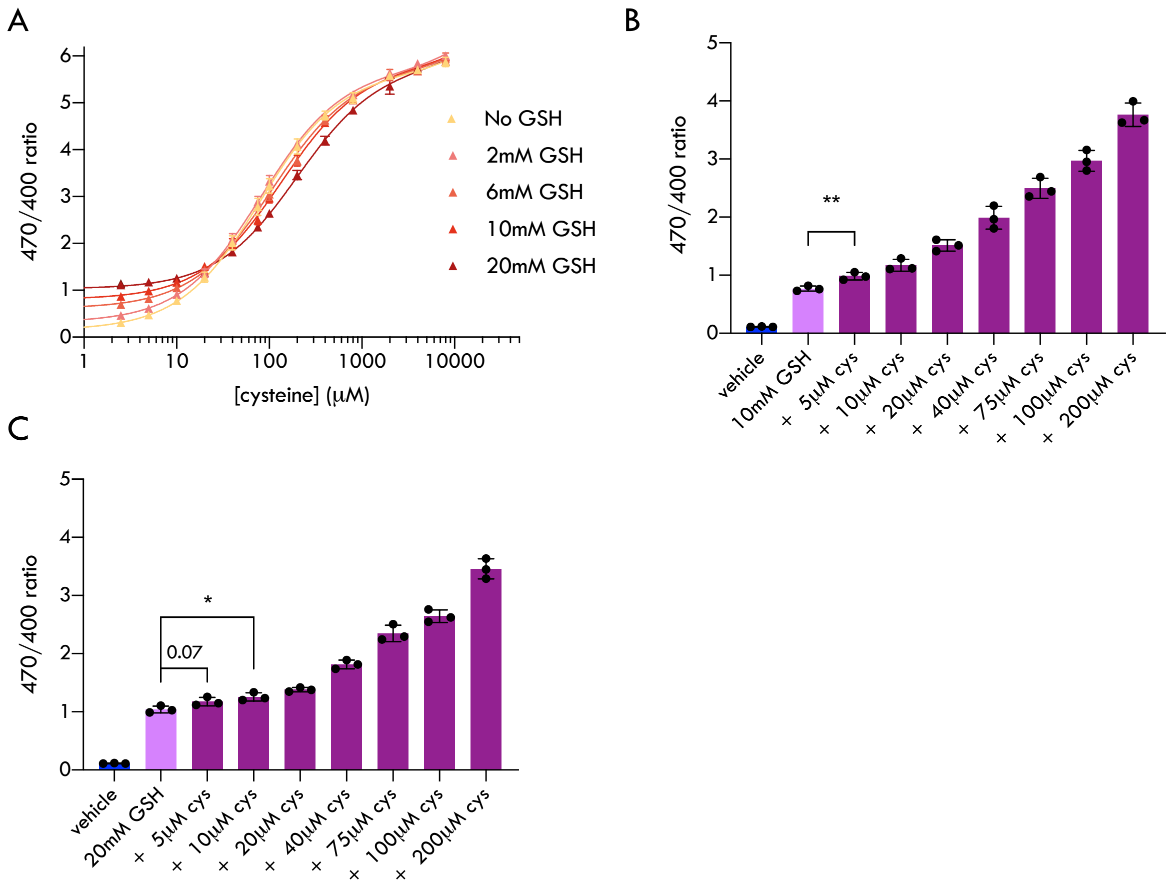

A
470/400 ratio
[cysteine] (μM)
No GSH
2mM GSH
6mM GSH
10mM GSH
20mM GSH
B
470/400 ratio
**
vehicle
10mM GSH
+ 5μM cys
+ 10μM cys
+ 20μM cys
+ 40μM cys
+ 75μM cys
+ 100μM cys
+ 200μM cys
C
470/400 ratio
*
0.07
vehicle
20mM GSH
+ 5μM cys
+ 10μM cys
+ 20μM cys
+ 40μM cys
+ 75μM cys
+ 100μM cys
+ 200μM cys

**Figure S3**

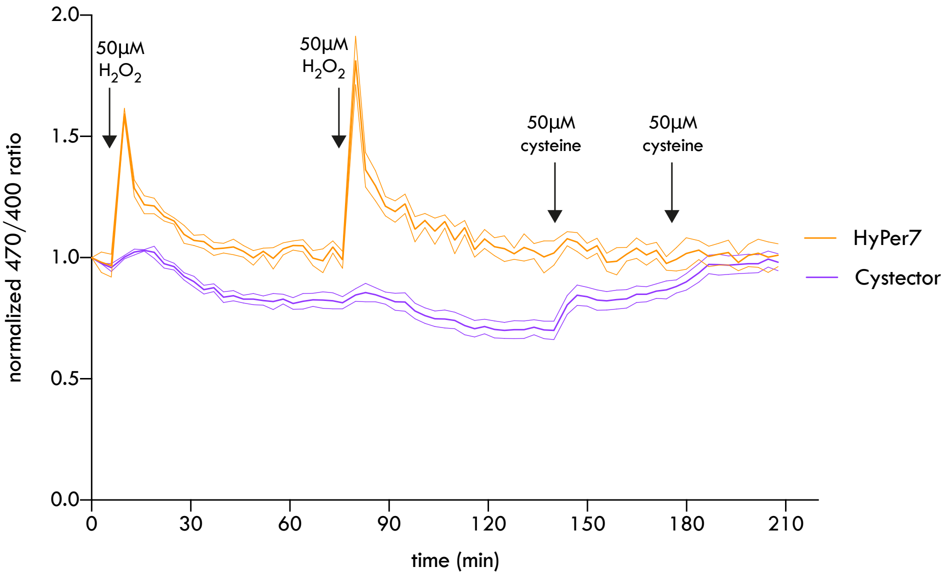

2.0
1.5
1.0
0.5
0.0
normalized 470/400 ratio
0
30
60
90
120
150
180
210
time (min)
50µM
$H_2O_2$
50µM
$H_2O_2$
50µM
cysteine
50µM
cysteine
HyPer7
Cystector
